# Generalized conditions for coexistence of competing parasitoids on a shared host

**DOI:** 10.1101/2020.12.24.424343

**Authors:** Abhyudai Singh

## Abstract

Motivated by the univoltine life histories of insects residing in the temperate-regions of the world, there is a rich tradition of modeling arthropod host-parasitoid interactions using a discrete-time formalism. We introduce a general class of discrete-time models for capturing the population dynamics of two competing parasitoid species that attack the same vulnerable stage of the host species. These models are characterized by two density-dependent functions: an *escape response* defined by the fraction of hosts escaping parasitism; and a *competition response* defined by the fraction of parasitized hosts that develop into adult parasitoids of either species. Model analysis reveals remarkably simple stability conditions for the coexistence of competing parasitoids. More specifically, coexistence occurs, if and only if, the adult host density increases with host reproduction rate, and the log sensitivity of the competition response is less than half. The latter condition implies that any increase in the adult parasitoid density will result in a sufficiently slow increase in the fraction of parasitized hosts that develop into parasitoids of that type. We next consider a model motivated by differences in parasitism risk among individual hosts with risk from the two parasitoid species assumed to be independently distributed as per a Gamma distribution. In such models, the heterogeneity in host risk to each parasitoid is quantified by the corresponding Coefficient of Variation (CV). Our results show that parasitoid coexistence occurs for sufficiently large reproduction rate, if and only if, the sum of the inverse of the two CV squares is less than one. This result generalizes the “CV greater than one” rule that defined the stability for a single parasitoid-host system to a multi parasitoid-host community.

## I. Introduction

The interaction between a single parasitoid species with its host is formulated as a discrete-time model

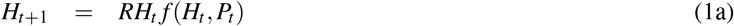

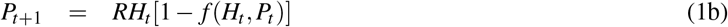

where *H_t_* and *P_t_* are the adult host, and the adult parasitoid densities, respectively, at the start of year *t* [1]–[3]. The model is motivated by a typical host/parasitoid life cycle as shown in Fig. 1, which consists of adult hosts emerging during spring, laying eggs that hatch into larvae [4]–[6]. Host larvae then overwinter in the pupal stage, and metamorphosize as adults the following year. Without loss of any generality, we assume that the host becomes vulnerable to parasitoid attacks in the larval stage. Adult female parasitoids emerge during spring, search and attack hosts by laying an egg into the body of the host.While adult parasitoids die after this time window, the parasitoid egg hatches into a juvenile parasitoid that grows at the host’s expense by using it as a food source, and this ultimately results in the death of the host. The juvenile parasitoids pupate, overwinter, and emerge as adult parasitoids the following year. In (1), *RH_t_* is the host larval density exposed to parasitoid attacks at the start of the vulnerable stage, where *R* > 1 denotes the number of viable eggs produced by each adult host. The function *f* (*H_t_, P_t_*) < 1 is the fraction of host larvae escaping parasitism and is referred to as the *escape response*. Thus, *RH_t_ f* (*H_t_, P_t_*) is the total larval density escaping parasitism to become adult hosts for next year. Finally, *RH_t_*[1 − *f* (*H_t_, P_t_*)] is the density of parasitized larvae that give rise to adult (female) parasitoids in the next generation.

**Fig. 1:**
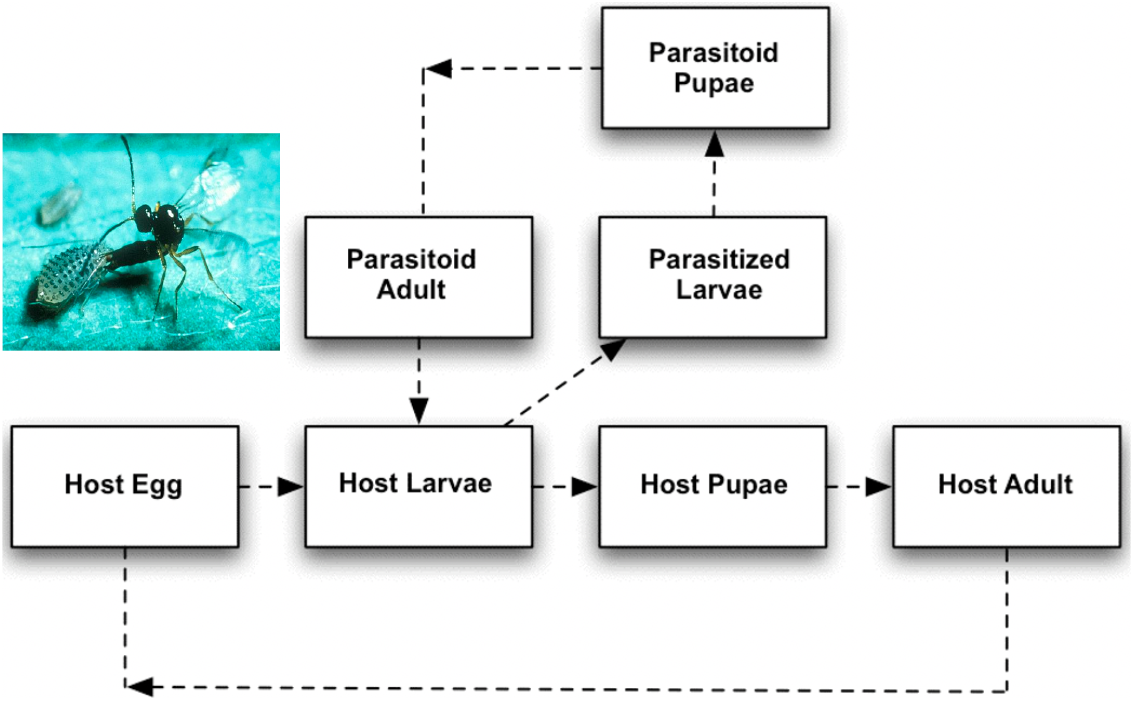
Life cycle of the host and the parasitoid. Inset shows the picture of a parasitoid wasp laying an egg into the body of its host (spotted alfalfa aphid). Picture taken from https://en.wikipedia.org/wiki/Parasitoid.

The simplest formulation of (1) is the classical Nicholson-Bailey model

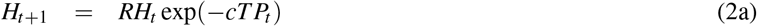

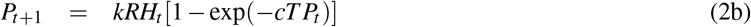

with a parasitoid-dependent escape response exp(−*cT P_t_*),where *c* > 0 represents the rate at which parasitoids attack hosts, and *T* is the duration of the host vulnerable stage [7]. The Nicholson-Bailey model is characterized by diverging oscillations in population densities resulting in an unstable population dynamics [7]. Recent work has identified two orthogonal mechanisms by which stability can arise in models of the form (1):

- The first mechanism is when the escape response *f* (*P_t_*) only depends on the parasitoid density, and then the non-trivial host-parasitoid equilibrium is stable, if and only, if, the equilibrium adult host density is an *increasing* function of the host reproduction rate *R* [8]. This type of stability arises through several related processes, such as, a fraction of the host population being in a refuge (i.e., protected from parasitoid attacks) [3], [9], large host-to-host difference in parasitism risk [8], [10]–[12], parasitoid interference [13]–[15], and aggregation in parasitoid attacks [16]–[18].
- The second mechanism is a Type III functional response where the parasitoid attack rate accelerates sufficiently rapidly with increasing host density [19]. Here the escape response *f* depends on both the host and parasitoid density, and interestingly, in this case stability leads to the adult host equilibrium density being a *decreasing* function of the host reproduction rate *R* [20].

A key focus of this work is to expand these results to multi-parasitoid communities. Towards that end, we consider two competing parasitoid species that attack the same vulnerable stage of the host species, as has been well documented in nature [21]–[27].

## II. Formulation of a parasitoid-competition model

Following along the lines of model (1), we introduce a novel class of parasitoid-competition models that take the form

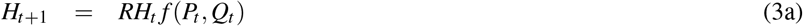

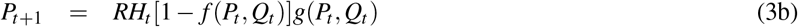

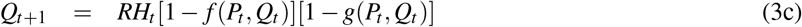

where *P_t_* and *Q_t_* represent the densities of the two competing parasitoids in year *t*. The escape response *f*(*P_t_, Q_t_*) is assumed to depend only on the parasitoid densities, and is a continuously differentiable function that is monotonically decreasing in both arguments. In the absence of both parasitoids

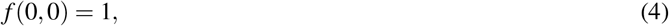

and the host population grows unboundedly as

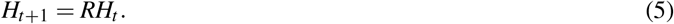

Recall that *RH_t_*[1 − *f*(*P_t_, Q_t_*)] in the net density of parasitized larvae. The function 0 ≤ *g*(*P_t_, Q_t_*) ≤ 1 represents the *competition response* that is the fraction of parasitized larvae that will develop into adult parasitoids *P*_*t*+1_ in the next generation. Similarly, 1 − *g*(*P_t_, Q_t_*) is the fraction of parasitized larvae that will develop into adult parasitoids *Q*_*t*+1_. To be ecologically relevant, *g*(*P_t_, Q_t_*) is an increasing function of *P_t_*, but a decreasing function of *Q_t_* with the following properties

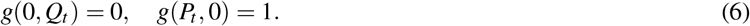

Generally, a semi-discrete framework is needed for a derivation of (3) that mechanistically captures the population interactions during the host’s vulnerable stage [**?**], [19], [28]–[31]. Taking a phenomenological approach, a simple example of the competition response is

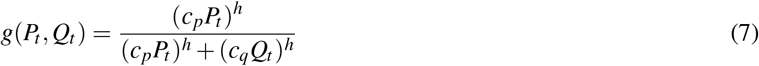

for positive constants *c_p_*, *c_q_* and *h* ≠ 1.

The non-trivial equilibrium densities *H*\*, *P*\*, *Q*\* of the competition model (3) satisfy

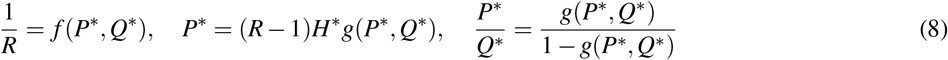

where the last equation determines the ratio of the parasitoid densities. For example, the competition response (7) leads to the ratio

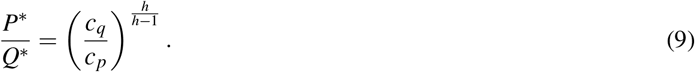

Before performing a systematic stability analysis, it is important to point out that model (3) generalizes previous multi parasitoid-host models, many of which implicitly assume that different parasitoid species attack different host developmental stages [32]–[36].

## III. Conditions for parasitoid coexistence

We begin by defining dimensionless log sensitivities of the equilibrium densities to the host reproduction rate *R*

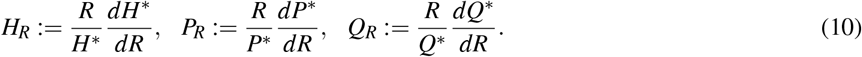

Similarly, we defining log sensitivities of the escape/competition response to the parasitoid densities

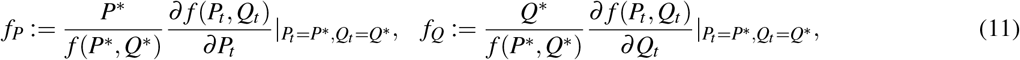

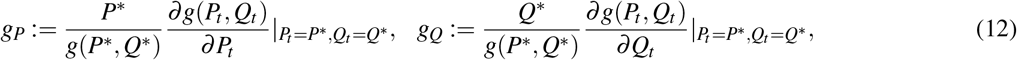

where 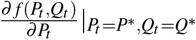 represents the partial derivative of the escape response with respect to *P_t_* evaluated at the equilibrium point. Note that for the competition response (7) the log sensitivities sum to zero, i.e.,

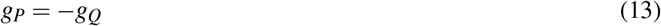

irrespective of *c_p_*, *c_q_* and *h*. Considering small perturbations *h_t_*, *p_t_* and *q_t_*

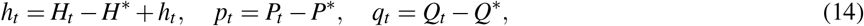

and linearizing model nonlinearities in (3) around the equilibrium, results in the following *linear* discrete-time system

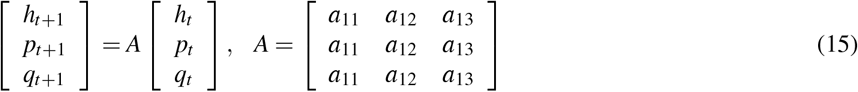

where the entries of the Jacobian matrix *A* are given by

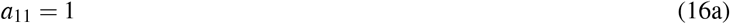

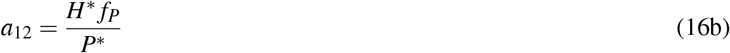

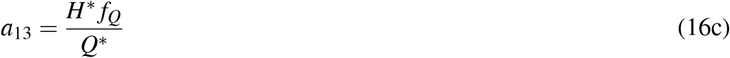

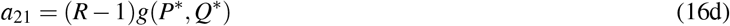

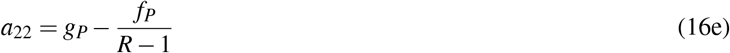

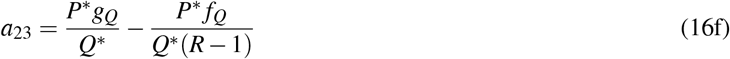

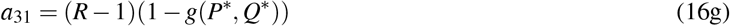

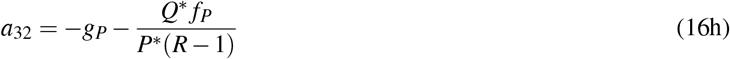

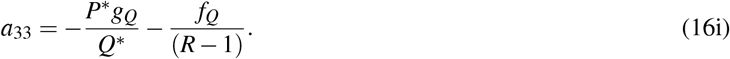

To derive analytical conditions for the stable coexistence of all three species we use the following result. Let

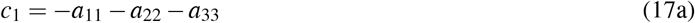

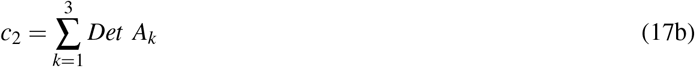

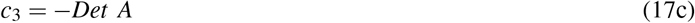

where *A_k_* is the 2 × 2 matrix obtained from matrix *A* by deleting row *k* and column *k*, and *Det* represents the matrix determinant. Then, the non-trivial fixed point *H*\*, *P*\*, *Q*\* is asymptotically stable, if and only if, the following inequalities hold

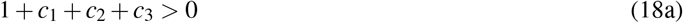

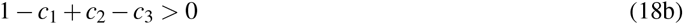

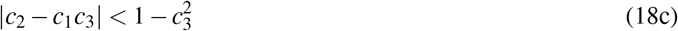

[37], [38].

### A. The symmetric case

Consider the symmetric case with equal densities *P*\* = *Q*\* and assuming (13), our analysis shows that inequality (18a) holds iff

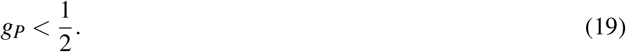

Inequality (18b) is always true and inequality (18c) holds iff

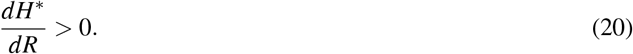

Thus, competing parasitoids can coexist as long as the adult host density is an increasing function of *R*, and a sufficiently low sensitivity of the competition response to the parasitoid density. For the symmetric case *c* = *c_p_* = *c_q_*, *P*\* = *Q*\*, 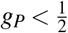 corresponds to having *h* < 1 in (7). While these results hold for any general function *f*, we illustrate them using the escape response

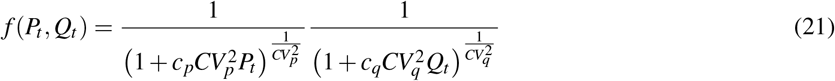

that is motivated by differences in parasitism risk among individual hosts. In particular, for each host, the parasitoids have a different attack rate (i.e., risk) that is assumed to be independent between the two species and follows a Gamma distribution [8], [12]. Here, *c_p_* and *c_q_* denote the average attack rates for parasitoids *P* and *Q* with the coefficient of variation in the attack rate given by *CV_p_* and *CV_q_*, respectively. An alternative interpretation of this escape response is that the parasitoid species independently aggregate attacks on a subpopulation of hosts with 1/*CV_p_* and 1/*CV_q_* quantifying the extent of clumping in parasitoid attacks [33]. For the symmetric case *c* = *c_p_* = *c_q_*, *CV* = *CV_p_* = *CV_q_*

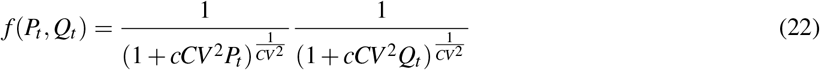

which solving (8) leads to the adult host equilibrium density

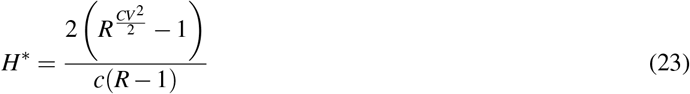

that is an increasing function of *R* (and hence, a stable equilibrium) iff *CV*^2^ > 2. In essence, coexistence of symmetric parasitoids on their shared host requires 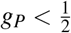 and *CV*^2^ > 2. Note that stable interaction of a single-parasitoid species and its host requires *CV*^2^ > 1 [8], [10]–[12], and our result *CV*^2^ > 2 shows that coexistence of multi-parasitoid communities requires hosts to have much larger variation in parasitism risk. These results are illustrated in Fig. 2 and 3 where a low sensitivity of the competition response (*h* < 1 in (7)) leads to coexistence of all three species. In contrast, a high sensitivity of the competition response (*h* > 1 in (7)) leads to extinction of one of the parasitoid species. It is interesting to point out that coexistence of competing parasitoids leads to a much lower host density in Fig. 2 as compared to a single parasitoid species in Fig. 3, and this has important implications for biological control of pests [39]–[42].

**Fig. 2:**
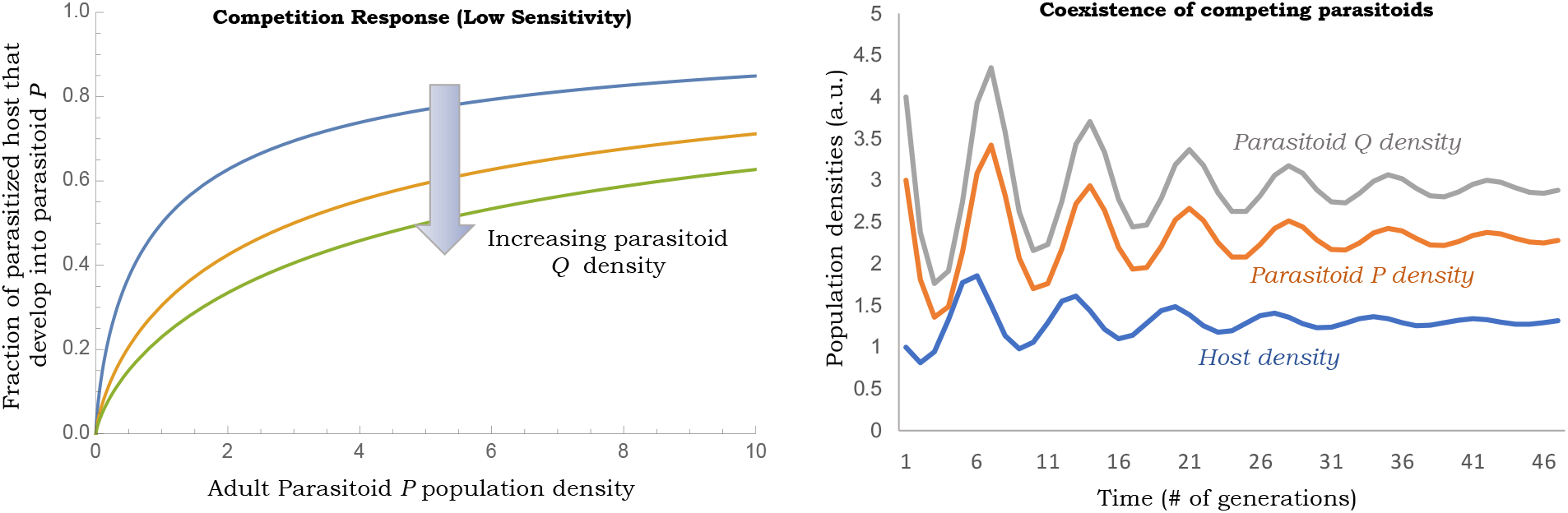
Low sensitivity of the competition response leads to coexistence of competing parasitoids together with their shared host. *Left*: Competition response (7) plotted as a function of parasitoid *P* population density for *h* = 0.7 and *c_p_* = *c_q_* = 1. *Right*: Simulation of model (3) with escape response (21) and competition response (7) for *c_p_* = 0.95, *c_q_* = 1.05, *CV* = *CV_p_* = *CV_q_* = 2.5. To contrast the population time series, parasitoid *Q* is assumed to have a slightly higher attack rate than parasitoid *P*.

**Fig. 3:**
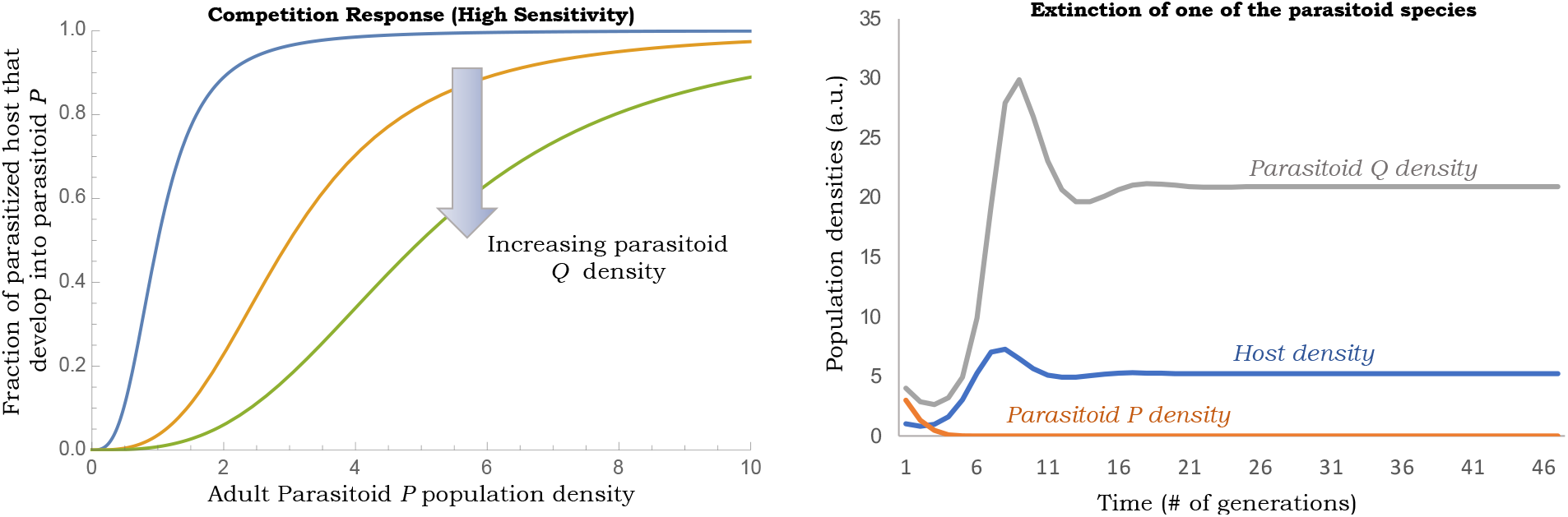
High sensitivity of the competition response leads to extinction of one the parasitoid species. *Left*: Competition response (7) plotted as a function of parasitoid *P* population density for *h* = 1.3 and *c_p_* = *c_q_* = 1. *Right*: Simulation of model (3) with escape response (21) and competition response (7) for *c_p_* = 0.95, *c_q_* = 1.05, *CV* = *CV_p_* = *CV_q_* = 2.5. The lower attack rate of parasitoid *P* results in its extinction, in spite of having *CV* > 2.

### B. The asymmetric case

We next turn our attention to the asymmetric case where *g*(*P*\**, Q*\*) ≠ 1/2 that results in

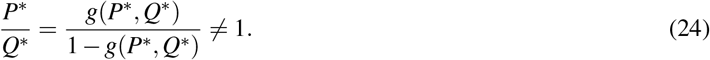

It turns out that assuming (13), inequality (18a) becomes

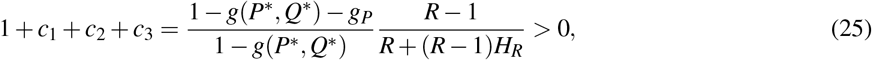

resulting in the stability condition

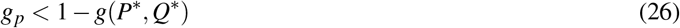

together with (20) that is still needed to satisfy inequality (18c). A simple interpretation of (26) is that as one parasitoid species gets more dominant, coexistence requires the competition response to become even more insensitive. Revisiting the escape response (21) and assuming *R* ≫ 1 such that

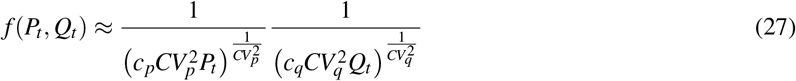

results in

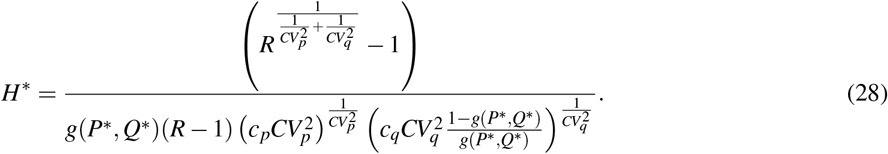

The adult host density being an increasing function of *R* leads to the stability criterion

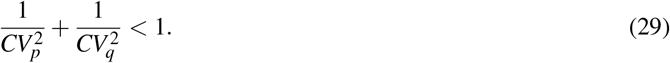

For the symmetric case *CV* = *CV_p_* = *CV_q_*, (29) reduces to our earlier result *CV*^2^ > 2, and also implies that having both 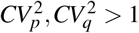 is a necessary condition for coexistence.

In summary, we have introduced a class of discrete-time models that captures competition of parasitoids on a shared host. The necessary and suffiicient conditions for coexistence of all three species turns to be simple and elegant: the adult host equilibrium increases with the host reproduction rate and the inter-species competition is sufficiently small. The latter condition is captured by the log-sensitivity of the competition response being low as given by (26) where any increase in the parasitoid *P* density does not lead to a sharp decrease in the parasitized larvae for parasitoid *Q*. Our stability results can easily be tested with field observation by monitoring population densities across generations. Future work will focus on incorporating correlations in the attack rate, allowing for one of the parasitoid to have a Type III functional response (which can also be strongly stabilizing [19]), and also exploring spatial mechanism for parasitoid coexistence [43]–[46].

